# Preserving Tissue Integrity Under the Beam: High-Energy, Low-Dose Synchrotron CT for *in situ* Imaging of Bovine Intervertebral Discs

**DOI:** 10.1101/2025.10.14.682281

**Authors:** C.M. Disney, R. Lees, A.W. Parker, R. Atwood, G. Burca

## Abstract

*In situ* tomography enables non-destructive, time-lapse imaging of biological tissues under load, offering insights into structural and mechanical changes. However, repeated scans can expose samples to high radiation doses, potentially altering tissue properties. This study evaluated the feasibility of low-dose synchrotron computed tomography (sCT) for high-resolution, *in situ* imaging of intact bovine intervertebral discs (IVDs), and assessed the effects of repeated X-ray exposure on mechanical, microstructural, and molecular integrity.

Intact oxtail IVD segments were imaged using propagation-based phase contrast sCT at 54 keV. Scan parameters were optimised to achieve high image quality within 66 seconds per scan, resulting in a total dose of ~30 kGy over six scans. Mechanical properties were assessed under cyclic loading, microstructural changes via digital volume correlation (DVC), and molecular alterations using Raman spectroscopy.

High-resolution imaging of soft and calcified tissues was achieved. Changes in sample stiffness, hysteresis, or stress recovery between irradiated and control were not identified. DVC revealed no microstructural damage or strain accumulation in the calcified endplate. Raman spectroscopy indicated minimal changes in soft tissues, with bone showing slight increased collagen crosslinking and reduced mineralisation.

Overall, this study demonstrates that high-energy, low-dose sCT enables repeated imaging of musculoskeletal tissues without compromising integrity, supporting its application in dynamic, time-lapse imaging studies. Importantly, larger, intact samples—such as whole bovine IVDs— were imaged overcoming limitations of previous studies that relied on small animal models. This approach supports more physiologically relevant investigations of tissue mechanics and degeneration in complex systems.

## 1. Introduction

MicroCT is a powerful three-dimensional imaging technique used to characterise whole biological tissues at the micro-to nanoscale. Compared to other imaging modalities, microCT offers the advantage of imaging large samples at high resolution. It has been most widely applied to bone and other calcified tissues, due to the absorption of X-rays by heavier elements such as calcium to generate contrast between structures. More recently, interest has grown in imaging soft tissues, using either staining techniques [1] or phase contrast methods [2]. While staining has proven effective for certain anatomical studies, it presents limitations, including the need for prolonged fixation times and potential alterations to tissue structure and mechanical properties [3]. In contrast, phase contrast imaging does not require staining; instead, it exploits differences in tissue refractive index. The simplest form of phase contrast imaging is propagation-based (or in-line), which requires only a semi-coherent X-ray source and a detector positioned at a distance downstream from the sample. This propagation distance causes Fresnel fringes to appear at the edges of structures, enhancing image contrast.

Samples can also be imaged under varying conditions or over time. Biomechanical *in situ* microCT studies of musculoskeletal tissues involve the application of mechanical loads to tissues or whole joints, while simultaneously resolving microstructural changes under near-physiological conditions [4]. Short scan times are particularly desirable for soft tissue samples, which exhibit long relaxation times, to minimise movement artefacts during imaging. Laboratory-based microCT systems are generally limited by longer scan durations and a trade-off between resolution and available field-of-view. In contrast, synchrotron CT (sCT) utilises larger, high-flux beams, enabling the imaging of larger samples at high resolution within significantly shorter scan times (ranging from seconds to minutes). However, a key limitation of *in situ* microCT studies is the requirement for multiple scans, which can expose samples to high levels of ionising X-ray radiation. Given the growing potential of sCT imaging and processing for biomechanical applications [5], there is now a pressing need to collectively focus on reducing X-ray exposure and understanding the effects of radiation on biological tissues during such experiments.

X-ray radiation can cause protein damage and cell death through high-energy photon ionisation processes such as absorption and scattering. The extent of radiation damage depends on several factors, including sample characteristics (composition, structure, and environmental conditions), beam properties (energy and flux), and imaging acquisition parameters (such as time in the beam, which is determined by projection exposure time and the number of projections). Barth et al. (2011) investigated the effects of radiation dose on cortical bone, demonstrating that collagen damage can impact tissue properties across multiple length scales. At the macroscale, irradiated samples became brittle, and microstructural cracks were visible in the image data following prolonged X-ray exposure.

These changes were attributed to reduced collagen fibrillar sliding due to crosslinking, as well as molecular damage to the primary structure of collagen. This seminal study by Barth et al. (2011) identified a dose of 35 kGy as a threshold often cited as a “safe” irradiation level—one that does not significantly compromise the mechanical integrity of cortical bone [6–9].

Previous attempts to limit x-ray damage have included optimising short scan times [10], controlling hydration conditions of the sample [11], and by filtering the x-ray beam [8]. Dose reduction through shorter scan times—achieved by decreasing either the number of projections or the exposure time per projection—has been explored extensively in bone *in situ* studies, but only more recently in soft tissue imaging [12]. Shorter scan times are more attainable for high contrast attenuating structures such as bone. However, soft tissues, resolved by phase contrast imaging have lower signal-to-noise and so require summation of signal leading to longer exposure times and higher number of projections.

Hydrated samples have been suggested to be more protected by water molecules from direct ionising protein damage [13]. Nevertheless, secondary damage may still occur through the ionisation of water molecules, which generates reactive species such as hydroxyl (OH) radicals that can damage proteins. Lowering the temperature of samples has also been investigated as a strategy to mitigate against direct and secondary damage [11]. Additionally, when using a polychromatic beam, filtering out lower-energy photons has been used to reduce potential damage caused by their higher absorption rates [7, 8, 14].

Methods of assessing x-ray damage include mechanical testing of samples, identifying microstructural changes of the sample in repeat scans [4, 7], and correlative molecular-scale assessment using techniques such as multiphoton microscopy [13], small angle x-ray scattering or x-ray diffraction, and Raman spectroscopy [15]. Mechanical testing of samples has shown that samples become stiffer and more brittle with increasing x-ray exposure [6, 7, 15, 16]. Digital volume correlation (DVC) can measure sub-voxel displacements between image volumes, and has been used to quantify microstructural changes such as microcrack and strain development in bone due to sample irradiation from multiple repeat scans [4, 7, 16, 17].

At the molecular level, multiphoton microscopy has revealed a loss of collagen triple helix structure, indicated by a reduction in second harmonic generation signal (Sauer et al., 2022). X-ray diffraction studies have reported strain relaxation in apatite crystals [13, 15], while Raman spectroscopy has identified increased collagen crosslinking [15]. Notably, these investigations have primarily focused on calcified tissues, and the potential for radiation-induced damage in soft tissues remains underexplored.

In this study, we aim to optimise low-dose sCT of intervertebral disc (IVD) oxtail samples and to quantify potential changes in their mechanical properties, microstructural integrity, and molecular changes due to x-ray irradiation of multiple scans.

## 2. Methods

### 2.1. Sample preparation

Oxtail segments were obtained from a local abattoir (Vicars game, Reading UK). A band saw was used to cut through adjacent vertebra (second-third and third-forth coccygeal vertebrae), leaving the disc intact. Excess muscle tissue was removed from each segment, the samples wrapped in PBS-soaked tissue paper, then frozen and stored at −80°C.

### 2.2. sCT imaging

Synchrotron tomography was conducted at the I12-JEEP beamline of Diamond Light Source [18] using a 54 keV monochromatic beam. A scintillator coupled with a PCO Edge camera and optical magnification provided an effective pixel size of 3.24 μm across an 8 × 7 mm field of view.

To investigate image quality, three different projection exposure times were selected: 0.018 s (minimum), 0.024 s (intermediate), and 0.030 s (maximum). These were determined based on the maximum exposure achievable without oversaturating the flat field, both with and without the Deben mechanical testing rig in place. The rig’s glassy carbon tube (6 mm in the beam path) acted as a filter, increasing the required exposure time by a factor of 1.66.

The number of projections collected over 180° sample rotation varied between 600 to 4000. The detector was positioned 2 m away from the sample to allow for propagation-based phase contrast image of the soft tissues [19].

Image volumes were reconstructed using the Savu pipeline at Diamond Light Source[20]. Flat- and dark-field corrected projections underwent ring artefact removal[21]. A Fresnel filter (value: 450) was applied to enhance contrast in intervertebral soft tissues. Reconstruction was performed using filtered back projection, followed by down-sampling (mean binning, bin size = 3 × 3 × 3) and rescaling to 8-bit using 2% saturation of the stack histogram.

Image quality was assessed using the middle 200 slices with the Perception based Image Quality Evaluator (MATLAB version: 9.13.0 (R2022b), Natick, Massachusetts: The MathWorks Inc.; 2022), which provides a no-reference quality score [22].

### 2.3. Dose calculation

Radiation dose was calculated separately for intervertebral disc and vertebral bone tissues. Elemental composition and density values for each tissue type were obtained from the IT’IS database [23]. Beam flux was determined after accounting for attenuation by the Deben rig’s glassy carbon support tube, which had a wall thickness of 3 mm.

Using the XCOM: Photon Cross Sections Database [24], mass attenuation coefficients—including contributions from coherent scattering—were calculated at 54 keV for each tissue based on their elemental composition. The absorbed flux and corresponding absorbed energy were then derived from these coefficients. Finally, dose rate was calculated by incorporating the respective tissue densities.

### *2*.*4. In situ* mechanical testing strategy

Samples were mechanically tested using a Deben CT 5000 rig equipped with a 5 kN load cell. The rig allows for up to 10 mm of extension with a resolution of 300 nm. To accommodate samples ranging from 20–30 mm in height, the support tube was extended using machined aluminium ring spacers. Samples were mounted using super glue onto custom 3D-printed holders designed to fit the Deben jaws.

The testing protocol was based on Newell et al 2020 [25], included an equilibration period to preload the sample equivalent to the intradiscal pressure of a healthy individual during lying postures, and force-controlled cyclic loading. During equilibration, a preload of 50 N was applied for 6 minutes, followed by a 2-minute recovery at 5 N, and then another 6-minute hold at 50 N. Cyclic loading was performed between 50 N and 800 N at a motor speed of 1 mm/min. Preliminary testing indicated that four cycles were sufficient to achieve a repeatable force-displacement response.

After each set of cyclic loading and the force returning to 50 N, the rig position was held for 2 minutes and then the sample scanned. This was repeated for 6 sets of cyclic loading and scans. Control samples underwent the same mechanical loading protocol but were not exposed to X-ray irradiation.

Measurements were taken from the final cycle using custom MATLAB scripts (version: 9.13.0 (R2022b), Natick, Massachusetts: The MathWorks Inc.; 2022). Compression stiffness was calculated from the linear region of the force-extension curve between 600 N and 800 N. Hysteresis was quantified as the area enclosed by the force-extension loop. Recovery behaviour following cyclic loading was characterised by fitting the force-extension data to a double exponential model:

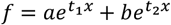

Where f is force, a and b are coefficients, t1 and t2 are time constants, and x is extension.

### 2.5. Quantifying microstructural changes in the mineralised endplate using digital volume correlation

Digital volume correlation (DVC) was used to assess microstructural changes in the calcified endplate due to the repeat scans. A point cloud specified the locations of each correlation point. Briefly, point clouds were created using a combination of image processing using Avizo and MATLAB: i) Calcified structures were segmented, ii) a finite element mesh created, iii) the endplate surface was selected as boundary nodes, iv) boundary nodes were exported to MATLAB for a surface fit to create a dense, evenly spaced point cloud which extended into the thickness of the endplate, v) the point cloud was imported back into Avizo to exclude points which lie in large pores. For DVC, spherical sub-volumes with 25-pixel diameters were chosen after optimising their size (Supplementary figure 2). DVC was run and Lagrange strains calculated using the Collaborative Computational Project in Tomographic Imaging (CCPi) iDVC app (https://tomographicimaging.github.io/iDVC, last accessed June 2025).

### 2.6. Quantifying molecular changes using Raman microscopy

For Raman microscopy, a Renishaw inVia Reflex Raman Microscope was used equipped with a Renishaw High Power Near infrared 830 nm diode laser with maximum output of 55 mW. Firstly, the sample was mounted as flat as possible inside a glass-bottomed dish with the glass insert removed (prepared previously using ethanol/methanol to dissolve the glue). This left an opening in the plastic dish where the upright Raman microscope objective could access the sample without the presence of glass contaminating the signal. The sample was kept moist with paper towel soaked in PBS surrounding it. The area of interest was identified, and a line scan of 1 mm steps was produced from the centre to the edge of the intervertebral disc.

A 5x magnification objective (Leica N PLAN 5x/0.12) was used to obtain the images, with a 1200 lines/mm grating, 10 seconds exposure time, and multiple accumulations for signal to noise averaging (3 accumulations for bone, 15 accumulations for intervertebral disc). The spectral scan was completed across 300 to 1800 cm^−1^ with cosmic ray removal enabled and autofocus enabled to ensure the surface of the sample was kept in focus for the duration of the line scan. The laser power was set to 100% and an ND filter of 0.0005% was used to reduce power before the sample. Spectra were collected using a 600 g/mm grating and a cooled CCD. The two-dimensional spectral images were assembled using the dedicated software.

A Python script was written for spectral filtering and baseline removal in the range of 765 to 1720 cm^−1^. To enhance signal quality, a Savitzky-Golay filter (savgol filter from scipy.signal)[26] was applied to the data. This method smooths spectral data by fitting successive polynomial functions to minimise noise whilst preserving important features. A window length of 41 and polynomial degree of 8 was chosen for the filter. Zhang baseline removal (ZhangFit from BaselineRemoval) [27] was used as it does not require peak information prior to processing. Spectral data was normalised to the Phenylalanine band at 1003 cm^−1^ to account for measurement variations. Phenylalanine, an essential amino acid, was chosen since it provides a stable reference of overall protein content. A previous study by Crawford-Manning et al. [28] provided references to identify spectra peaks of bovine intervertebral disc. The outer three and inner three spectra from each line scan were averaged to represent annulus fibrosus (AF) and nucleus pulposus (NP) spectra for each sample. A t-test across the whole spectra was then performed to assess statistical significance between irradiated and control sample groups. Bone spectral data were available from only four samples (two irradiated and two control). Twenty-five bone spectra were averaged for each sample, and principal component analysis was applied to examine variance between spectra.

## 3. Results

### 3.1. Phase contrast imaging of bovine intervertebral disc resolves calcified endplate and collagen fibre bundle microstructure

Phase contrast imaging was used to resolve soft tissue microstructure within a whole bovine IVD. A representative local tomography scan of the annulus fibrosus–nucleus pulposus (AF–NP) boundary is shown in Figure 1. Virtual slices taken from the reconstructed image volume include a mid-transverse view (Figure 1a) and a sagittal view (Figure 1b). The mid-transverse slice reveals the concentric lamellar architecture of the AF, with lamellae decreasing in thickness towards the NP boundary (Figure 1a). The calcified endplates, located at the superior and inferior boundaries of the IVD, are clearly resolved in the sagittal slice through absorption contrast (Figure 1b).

**Figure 1.**
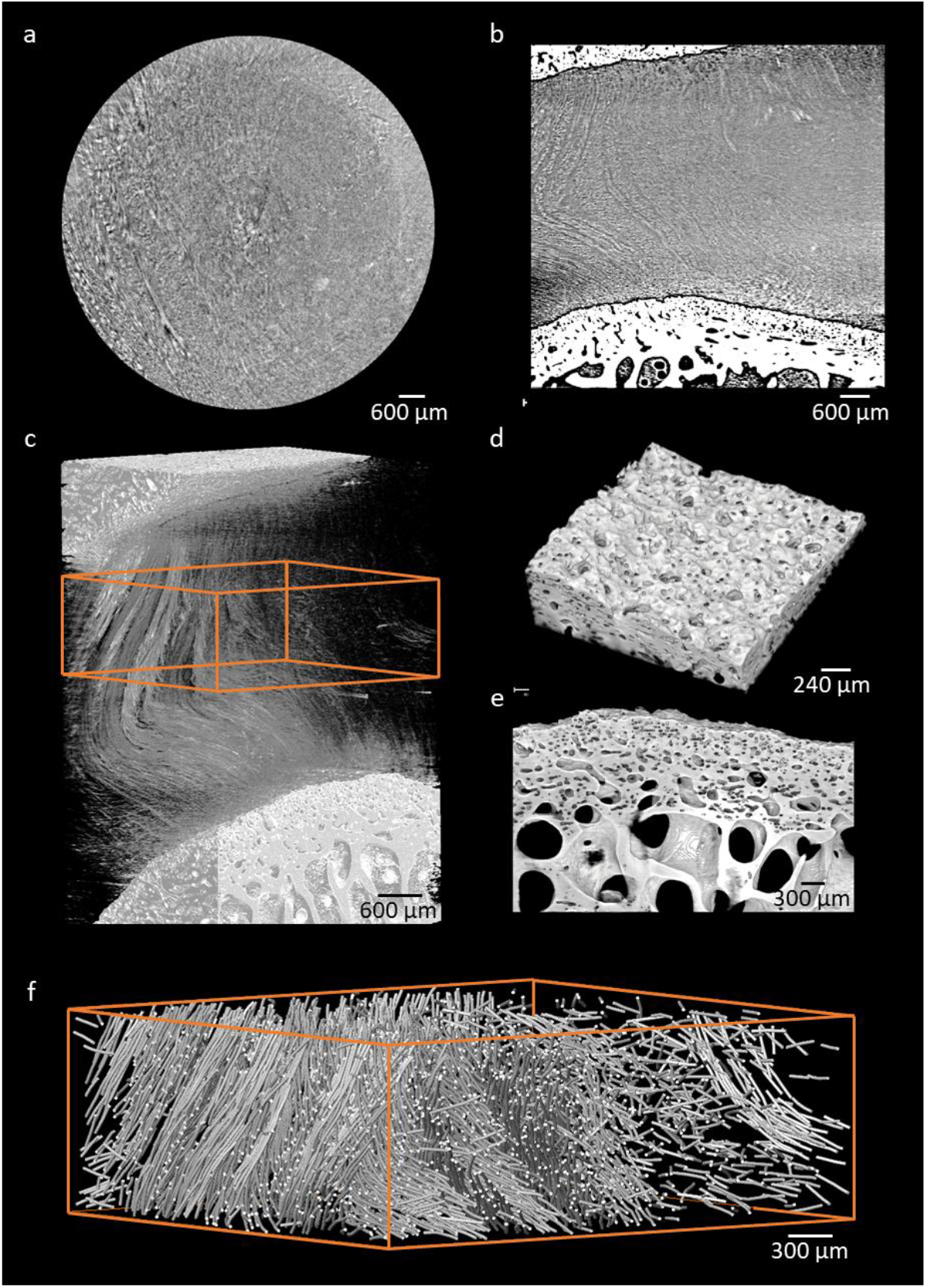
Phase contrast synchrotron tomography of a bovine intervertebral disc acquired at the I12 beamline, Diamond Light Source. a) Mid-transverse virtual slice showing concentric lamellae of the annulus fibrosus (AF). b) Mid-sagittal slice resolving both calcified endplates and soft tissue of the disc. c) Three-dimensional rendering of the AF collagen fibre bundles inserting into the calcified endplates. d & e) 3D renderings of the highly porous endplate structure. f) Traced individual collagen fibre bundles within a region of interest in the AF.

Three-dimensional renderings of the image volume further illustrate the AF lamellar microstructure, consisting of collagen fibre bundles that insert into the calcified endplate (Figure 1c). The highly porous endplate surface at the AF boundary (Figure 1d) was resolved connected to underlying trabecular architecture (Figure 1e). Individual collagen fibre bundles within the AF were resolved with sufficient contrast to trace using image processing (XFiber Avizo 3D extension [version 2021.1, Thermo Fisher Scientific]) (Figure 1f). Alternating fibre orientations between adjacent AF lamellae, as well as a progressive reduction in fibre density towards the NP boundary, could be observed.

To limit the time samples were exposed to the x-ray beam, exposure time per projection (0.03, 0.024 and 0.018) and number of projections per scan (4000 to 600) were tested for image quality in the AF. Image quality was quantified using the Perception-Based Image Quality Evaluator (PIQE), which detects image noise without requiring a reference image.

Representative mid-transverse slices from scans with the longest (180 s) and shortest (27 s) total scan durations are shown in Figure 2ai and 2bi, respectively. Corresponding PIQE noise maps are overlaid in Figure 2aii and 2bii, illustrating an increase in high-frequency noise with reduced scan time. PIQE score for the middle 200 slices for all scans is shown in Figure 2c, where an increase in PIQE score with shorter scan times indicates lower image quality. Notably, scan times below 60 seconds resulted in a marked degradation of image quality.

**Figure 2.**
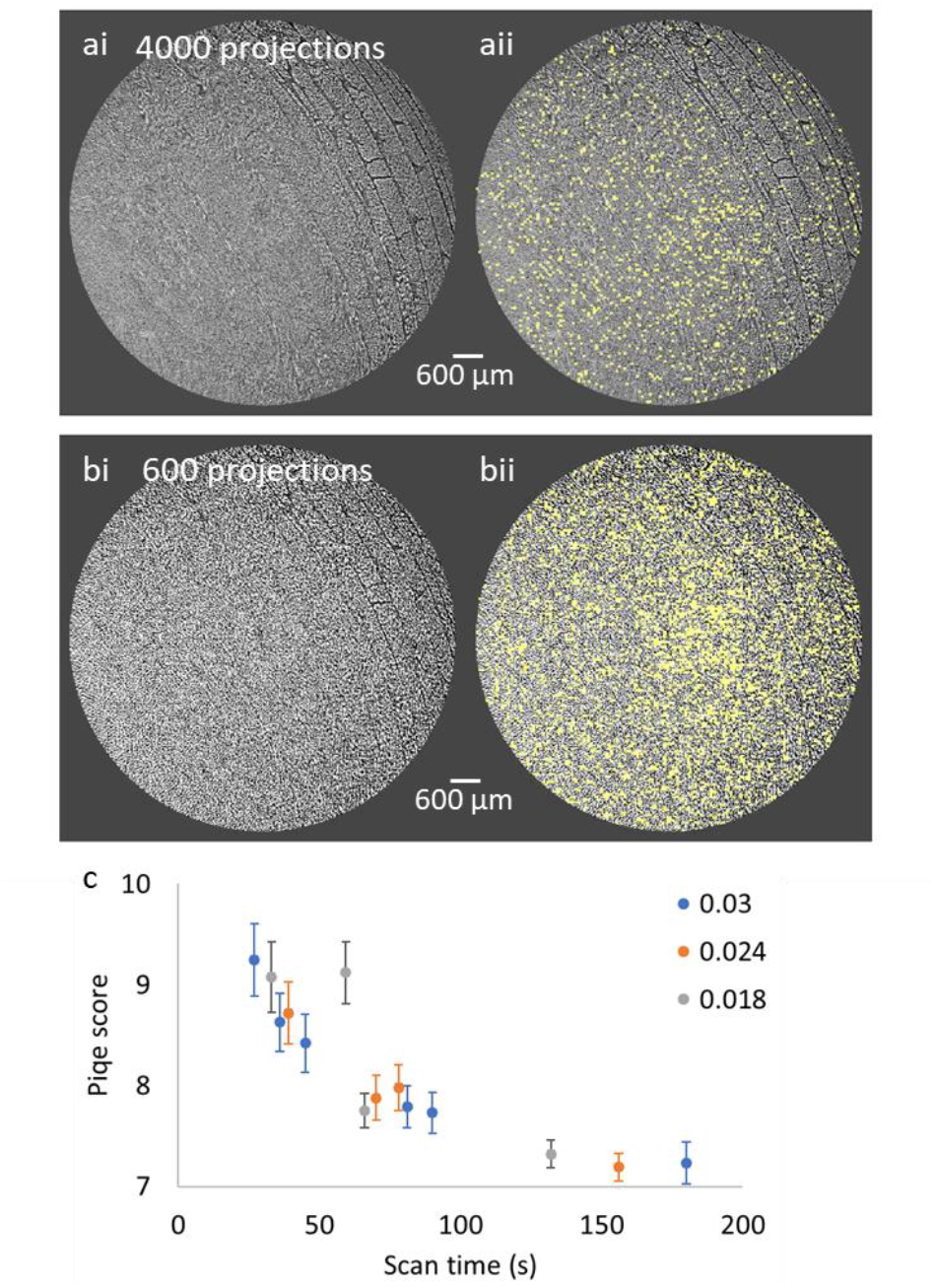
Optimisation of scan time and quantification of image quality. a) Example mid-transverse slice (i) and corresponding PIQE-detected noise overlay (ii) from a scan acquired with 4000 projections and a 0.030 s exposure time per projection. b) Example mid-transverse slice (i) and corresponding PIQE-detected noise overlay (ii) from a scan acquired with 600 projections and the same 0.030 s exposure time. c) Perception-Based Image Quality Evaluator (PIQE) scores for the central 200 slices across scans with varying projection exposure times (0.030, 0.024, and 0.018 s) and number of projections (4000 to 600), illustrating the relationship between scan duration and image quality.

X-ray absorption was 28% higher for the calcified tissues (vertebra) than the soft tissue (IVD) at 54 keV (Table 1). Dose rate, however, was slightly lower for vertebral tissues when compared to IVD due to its higher density (1 Gy = 1 J/kg).

**Table 1.** Absorbed flux energy and dose rate for IVD and vertebra tissues.

|  | IVD | Vertebra |
| --- | --- | --- |
| **Mass attenuation [cm<sup>2</sup>/g]** | 0.29 | 0.31 |
| **Tissue density [g/cm<sup>3</sup>]** | 1.10 | 1.91 |
| **Tissue thickness [cm]** | 2.50 | 2.00 |
| **Absorption** | 0.5464 | 0.6975 |
| **Absorbed energy flux [J/s/mm<sup>2</sup>]** | 0.0021 | 0.0027 |
| **Dose rate [Gy/s]** | 76.17 | 70.07 |

Time, dose and energy absorbed per scan is presented in Table 2 and Table 3. Based on the image quality (PIQE) scores in Figure 2, scans with 2000 projections at 0.018 s per projection were used for the remainder of the study which involved *in situ* mechanical testing over 6 scans. These scans had ~5 kGy dose and ~0.17 J/mm^2^ absorption per scan resulting in a total of ~30 kGy and ~1 J/mm^2^ for 6 scans. Note that tissue thickness is for the thickness part of the sample and assumed to be constant, meaning that doses are an over-estimate.

**Table 2.** Time per scan [s] including 0.015 s overhead per projection. Highlighted cell indicates the parameters used for the remainder of the study.

| **Projection exposure time [s]** | **Number of projections** |  |  |  |  |  |
| --- | --- | --- | --- | --- | --- | --- |
|  | **4000** | **2000** | **1800** | **1000** | **800** | **600** |
| **0.03** | 180.0 | 90.0 | 81.0 | 45.0 | 36.0 | 27.0 |
| **0.024** | 156.0 | 78.0 | 70.2 | 39.0 | - | - |
| **0.018** | 132.0 | 66.0 | 59.4 | 33.0 | - | - |

**Table 3.** Dose and absorbed energy per scan for IVD and vertebra tissues. Highlighted cell indicates the parameters used for the remainder of the study.

| Time per scan [s] | Dose per scan [kGy] |  | Absorbed energy per scan [J/mm<sup>2</sup>] |  |
| --- | --- | --- | --- | --- |
|  | IVD | Vertebra | IVD | Vertebra |
| 180 | 13.71 | 12.61 | 0.377 | 0.481 |
| 156 | 11.88 | 10.93 | 0.327 | 0.417 |
| 132 | 10.05 | 9.25 | 0.277 | 0.353 |
| 90 | 6.86 | 6.31 | 0.189 | 0.241 |
| 81 | 6.17 | 5.68 | 0.170 | 0.217 |
| 78 | 5.94 | 5.47 | 0.163 | 0.209 |
| 70.2 | 5.35 | 4.92 | 0.147 | 0.188 |
| 66 | 5.03 | 4.62 | 0.138 | 0.176 |
| 59.4 | 4.52 | 4.16 | 0.124 | 0.159 |
| 45 | 3.43 | 3.15 | 0.094 | 0.120 |
| 39 | 2.97 | 2.73 | 0.082 | 0.104 |
| 36 | 2.74 | 2.52 | 0.075 | 0.096 |
| 33 | 2.51 | 2.31 | 0.069 | 0.088 |
| 27 | 2.06 | 1.89 | 0.057 | 0.072 |

### 3.2. Sample mechanical properties of unirradiated and irradiated samples

*In situ* mechanical testing was performed between six scans to assess changes in whole sample mechanical behaviour (Figure 3). The testing protocol involved cyclic loading between 50 N and 800 N. A gradual accumulation of compressive creep strain in the soft tissue was observed across successive loading cycles. This is illustrated in Figure 3a, where the upward displacement of the endplate with each scan indicates progressive deformation.

**Figure 3.**
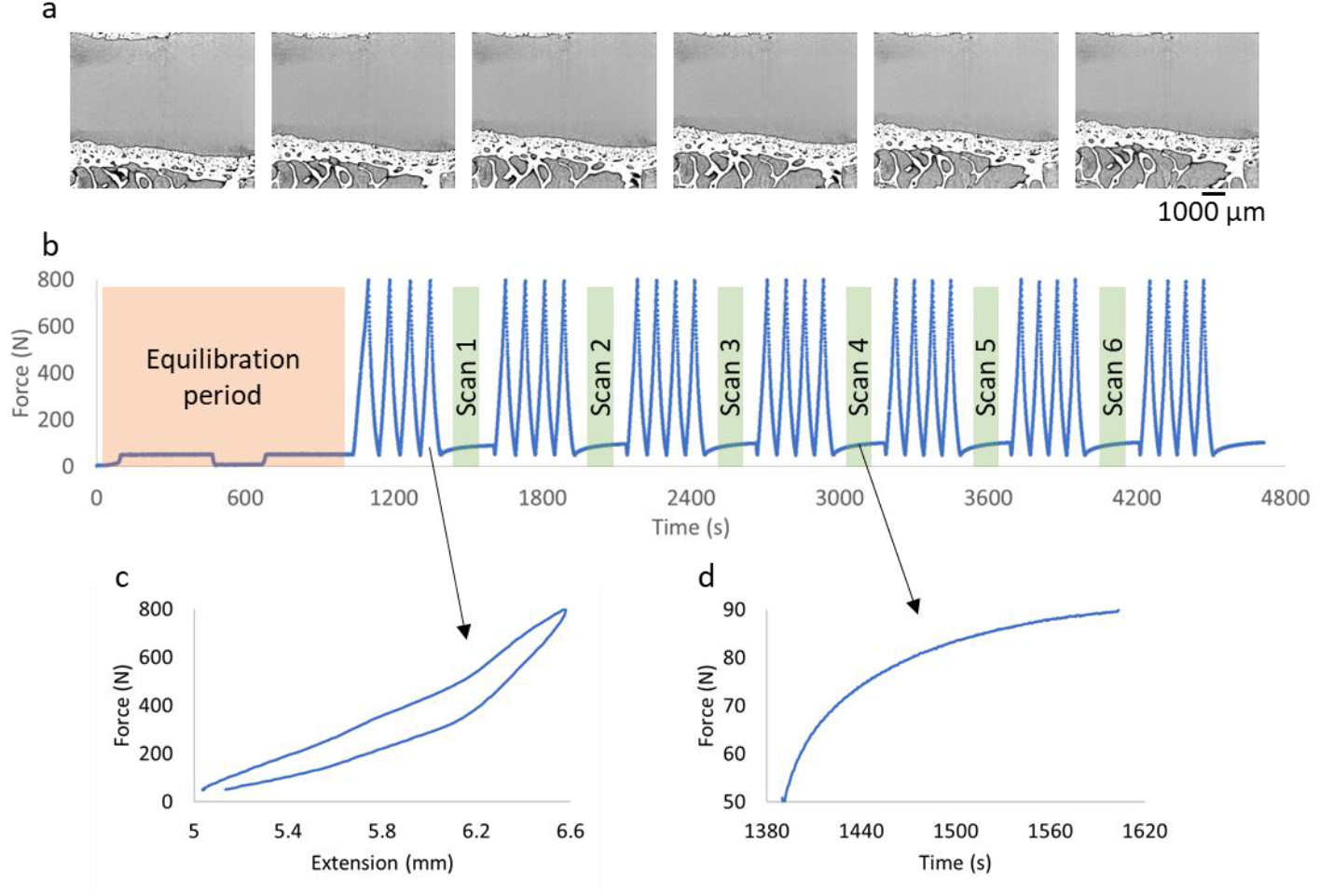
*In situ* mechanical testing set-up and response of the intervertebral disc during cyclic loading. a) Example mid-sagittal slices from each of the six scans, showing progressive upward displacement of the endplate due to compressive creep strain. b) mechanical testing strategy using force control. Initial low-level loading for sample equilibration, followed by cyclic loading between 50 N and 800 N, and a hold phase during scanning. c) Representative force–extension curve during cyclic loading, illustrating non-linear mechanical behaviour. d) Representative force– time curve showing stress recovery behaviour of the sample between loading cycles.

A consistent residual strain of approximately 20% was observed across all samples, as shown in Supplementary Figure 1.

Stiffness was calculated from the linear region of the force–extension curve between 600 and 800 N (Figure 4ai). All samples had an initial mean stiffness and standard deviation of 730 ± 142 N mm^−1^ and after the final cycle set 869 ± 177 N mm^−1^. The irradiated sample stiffness showed little variation with an increasing number of scans (703 ± 41, 921 ± 33, 558 ± 21 N mm^−1^) (solid lines in Figure 4 aii).

**Figure 4.**
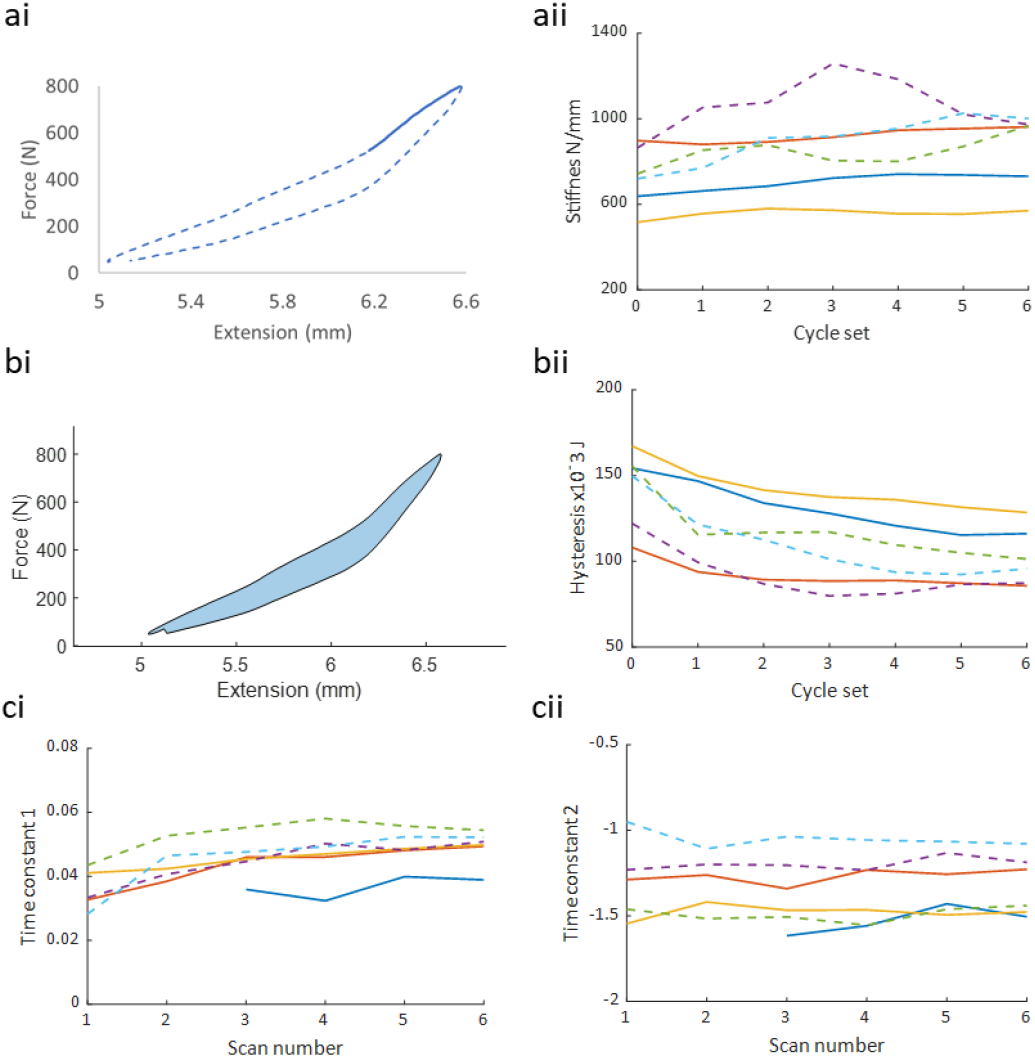
Whole-sample mechanical behaviour across cyclic loading and scanning. ai) Linear stiffness calculated from the final loading cycle between 600 and 800 N (highlighted by solid line). aii) Stiffness values across each cycle set. Unirradiated control samples are shown as dashed lines; irradiated samples as solid lines. The initial cycle set prior to the first scan is denoted as set 0. bi) Hysteresis quantified as the area enclosed by the force–extension curve. bii) Hysteresis values across cycle sets. Dashed lines represent unirradiated controls; solid lines represent irradiated samples. ci) and cii) Stress recovery time constants for unirradiated (dashed lines) and irradiated (solid lines) samples. Outliers from the first two scans of one unirradiated sample, caused by equipment error, were excluded.

Hysteresis, defined as the area enclosed within the force–extension curve, was used to quantify energy dissipation during each loading cycle (Figure 4bi). All samples demonstrated a gradual reduction in hysteresis with increasing cycle sets (Figure 4bii). Mean hysteresis values decreased from 142 ± 18 × 10^−3^ J to 102 ± 16 × 10^−3^ J between the first and final loading sets. The mean relative decrease in hysteresis was lower in irradiated samples (−23 ± 2%) compared to unirradiated samples (−33 ± 4%).

During the post-loading stress recovery phase, where samples were held at a constant position, force increased exponentially (Figure 3d). This recovery behaviour was characterised using time constants derived from an exponential model. Both irradiated and unirradiated samples exhibited similar recovery profiles (Figure 4c). Outliers caused by equipment error during the first two scans of one unirradiated sample were excluded from the analysis.

### 3.3. Microstructural integrity of mineralised endplate after multiple scans

DVC was used to identify any possible changes in calcified endplate microstructure or residual strain due to x-ray absorption after multiple scans. The highly porous architecture of the endplate provided a rich image texture, making it well-suited for DVC tracking. Sub-volume diameters ranging from 5 to 35 pixels were tested on 1000 points to determine optimal tracking performance (Supplementary Figure 2). A sub-volume diameter of 25 pixels (243 μm) demonstrated accurate tracking (median values ± 95^th^ percentiles; 0.0012 + 0.00266 for first principal strain and −0.0013 - 0.00271 for third principal strain) and was therefore used for subsequent analysis.

Over 112,000 points were placed throughout the calcified endplate across the full image volume (Figure 5a and b). Normalised sum-squared-difference (ZNSSD), used as the DVC minimisation objective function between sub-volumes, had a low narrow distribution and importantly showed no change with the increased number of scans (Figure 5c).

**Figure 5.**
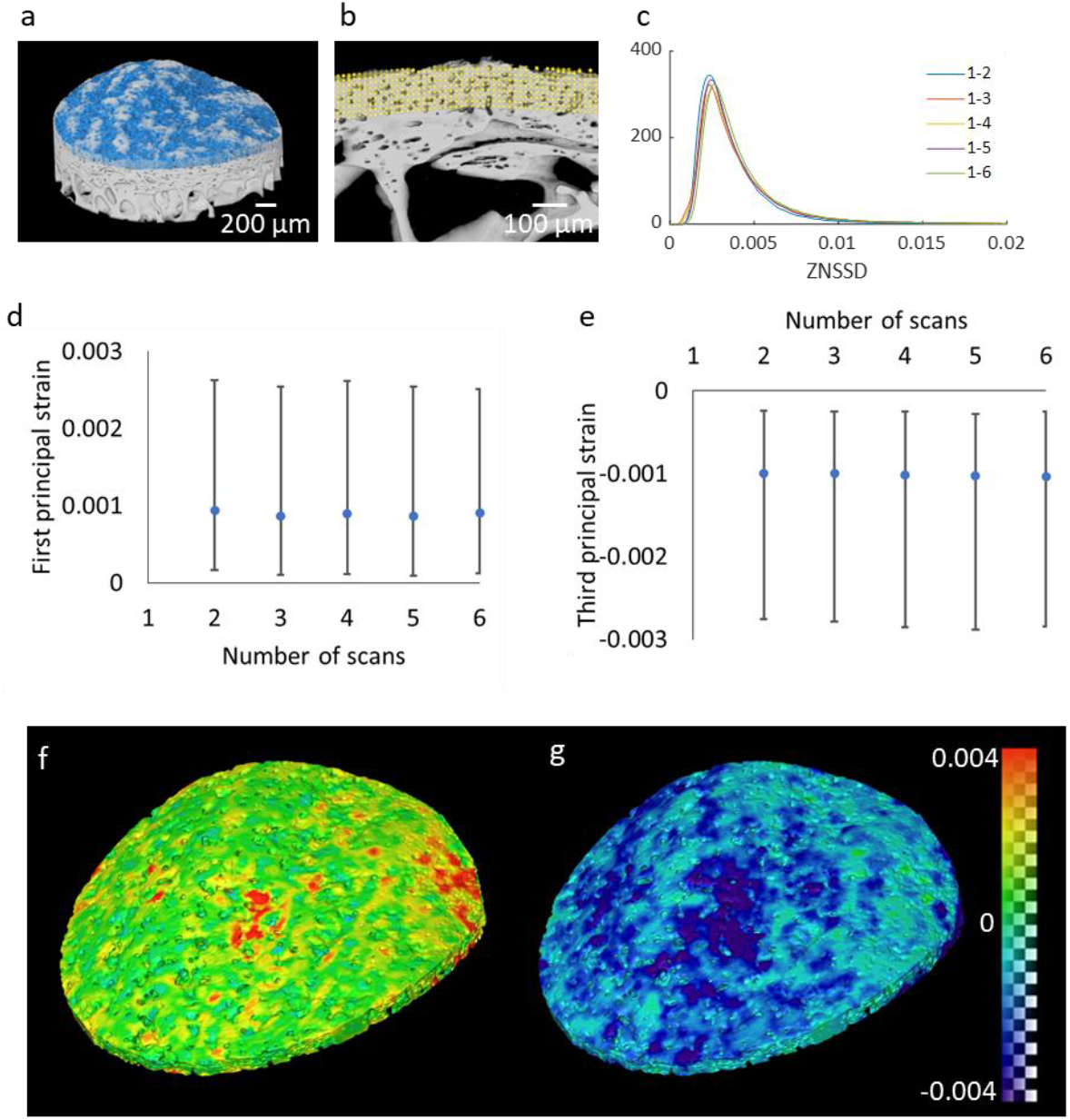
Assessment of microstructural changes and residual strain in the calcified endplate after multiple scans. a) Region of interest selected for DVC analysis, encompassing the entire calcified endplate (highlighted in blue in the 3D rendering). b) Dense point cloud spanning the full thickness of the endplate, with each point tracked across scans. c) DVC correlation residual (ZNSSSD: normalised sum square difference) between scans, where scan 1 is used as the reference. d) and e) Median values of first and third principal strains after each scan, calculated relative to the initial scan. Error bars represent the 5th and 95th percentiles. f) and g) Spatial maps of first and third principal strains between the first and sixth scans, overlaid on segmented image data.

First and third principal strains were used to find the maximum and minimum strain present at each measurement point. First and third principal strain had median values ~0.001 ε which remained constant across all 6 scans (Figure 5d and e). The variation in strain also remained constant with an increased number of scans as shown by the error bar at the 5^th^ and 95^th^ percentiles, without exceeding ±0.003 ε for both principal strains, suggesting no accumulation of strain due to repeated scanning.

The spatial distribution of strain values after the final sixth scan was mapped onto the segmented image data (Figure 5f and g). Strain remained low throughout the endplate, although some heterogeneous regions of elevated strain were observed. These areas, particularly near the centre, were associated with image artefacts such as central voids introduced during ring artefact removal (Supplementary figure 3).

### 3.4. Molecular changes measured by Raman

Raman spectroscopy was performed on samples following six X-ray tomography scans (30 kGy), as well as on unirradiated controls. Inspection of the AF Raman spectra (Figure 6a) showed no significant changes in collagen I-associated bands, including Amide I (1667 cm^−1^), Amide III (1245 and 1270 cm^−1^), and the collagen backbone (813 cm^−1^). However, small but statistically significant (p<0.05) changes were observed in the 1318 cm^−1^ (CH_3_, CH_2_, proline) and 1340 cm^−1^ (glycosaminoglycans, δ C– H (CH_2_)) bands. The NP spectra (Figure 6b) revealed minor but significant (p<0.05) shifts in the Amide I band (1667 cm^−1^). Bone spectra showed differences between irradiated and unirradiated samples (Figure 6c). The principal component analysis (PCA) loadings plot showed that there was an increase in collagen associated peaks (Amide I, CH2 bending mode, Amide III, collagen backbone), a decrease in phosphate peak, and shift in carbonate substitutions within the 1070–1100 cm^−1^ range.

**Figure 6.**
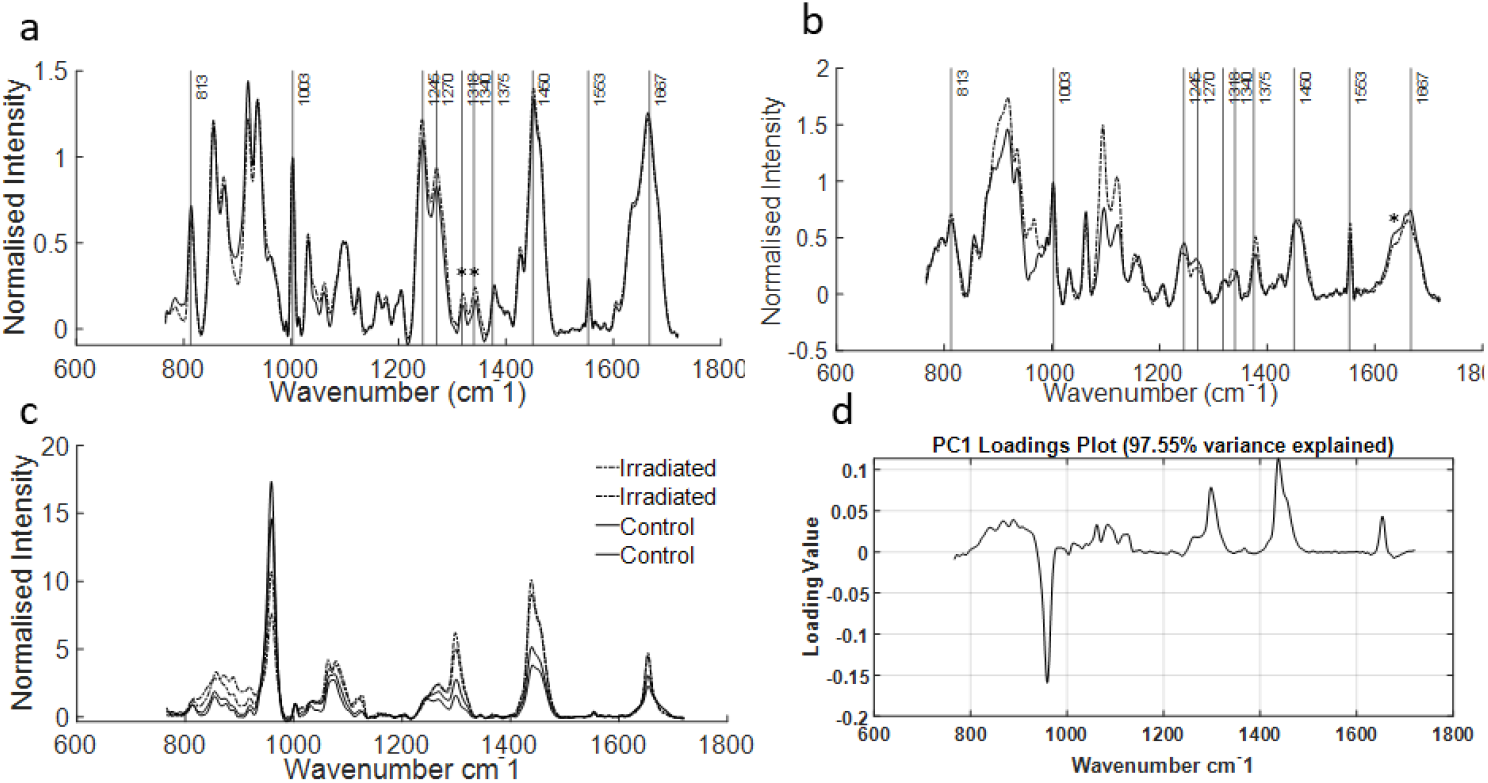
Detection of molecular changes in intervertebral disc and bone tissue using Raman spectroscopy. a) Averaged Raman spectra of the annulus fibrosus (AF) for irradiated (dashed line) and unirradiated (solid line) samples. b) Averaged spectra of the nucleus pulposus (NP) for irradiated (dashed line) and unirradiated (solid line) samples. Asterisks (*) indicate statistically significant differences (p < 0.05). c) Representative bone spectra from four samples (two irradiated and two unirradiated). d) Principal component analysis (PCA) loadings plot showing spectral variance between irradiated and unirradiated bone samples.

## 4. Discussion

### 4.1. Imaging of larger samples with shorter scan times

The capability to image larger samples is a crucial step closer to increasingly physiologically relevant human biomechanics and pathology studies. A major limitation of previous *in situ* musculoskeletal studies has been the requirement for small samples (on the millimetre scale) due to the beam size available. This has typically necessitated the use of small animal models or isolated segments of tissue.

In this study, we successfully imaged the microstructure of intact native bovine intervertebral disc from oxtails (Figure 1). The samples measured approximately 25 mm in diameter – substantially larger than those used in our previous work with lumbar rat spine (~5 mm diameter). Samples included vertebra, endplate and disc leaving the hard-to-soft tissue boundary intact which is crucial for maintaining tissue biomechanics and the main advantage of using sCT.

Despite this advancement, this study was still limited by the detector field-of-view meaning that only 8 by 7 mm local tomography scans within the sample was possible at 3.24 μm pixel size. However, emerging synchrotron technologies offer the potential for *in situ* imaging of larger samples with expanded fields of view at microscale resolution[5], paving the way for more comprehensive and translational biomechanical investigations.

In addition to the physical dimensions of the beam, sufficient X-ray transmission is essential for *in situ* imaging of large samples with centimetre-scale thickness. Adequate transmission is critical not only for achieving high-quality images but also for enabling short scan times, which help minimise motion artefacts during mechanical testing. In this study, a high-flux beam with higher energy (54 keV compared to 27 keV in previous work [Table 4]) was used to gain appropriate transmission through the sample whist achieving scans ~ 1 minute.

**Table 4.**
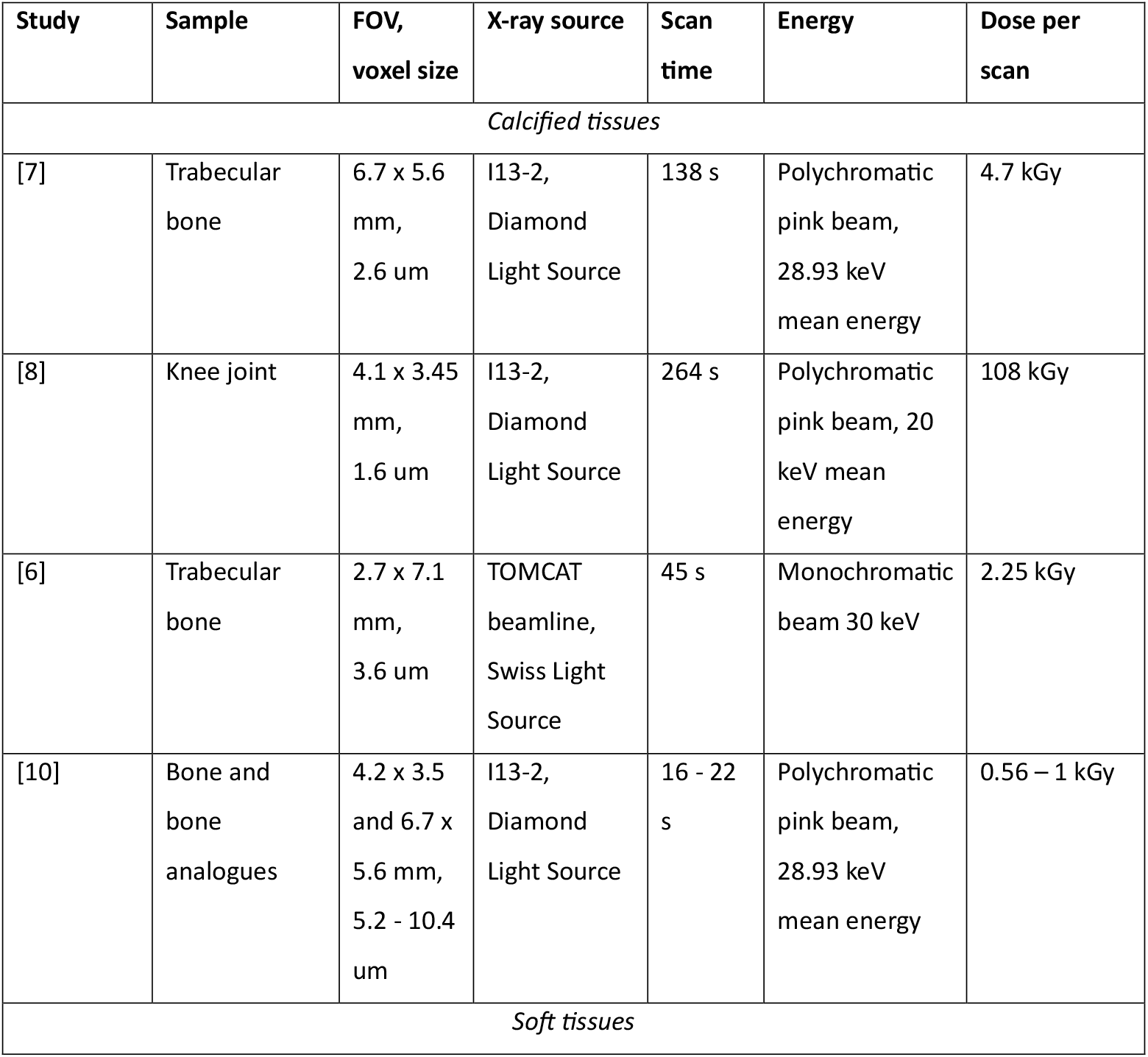
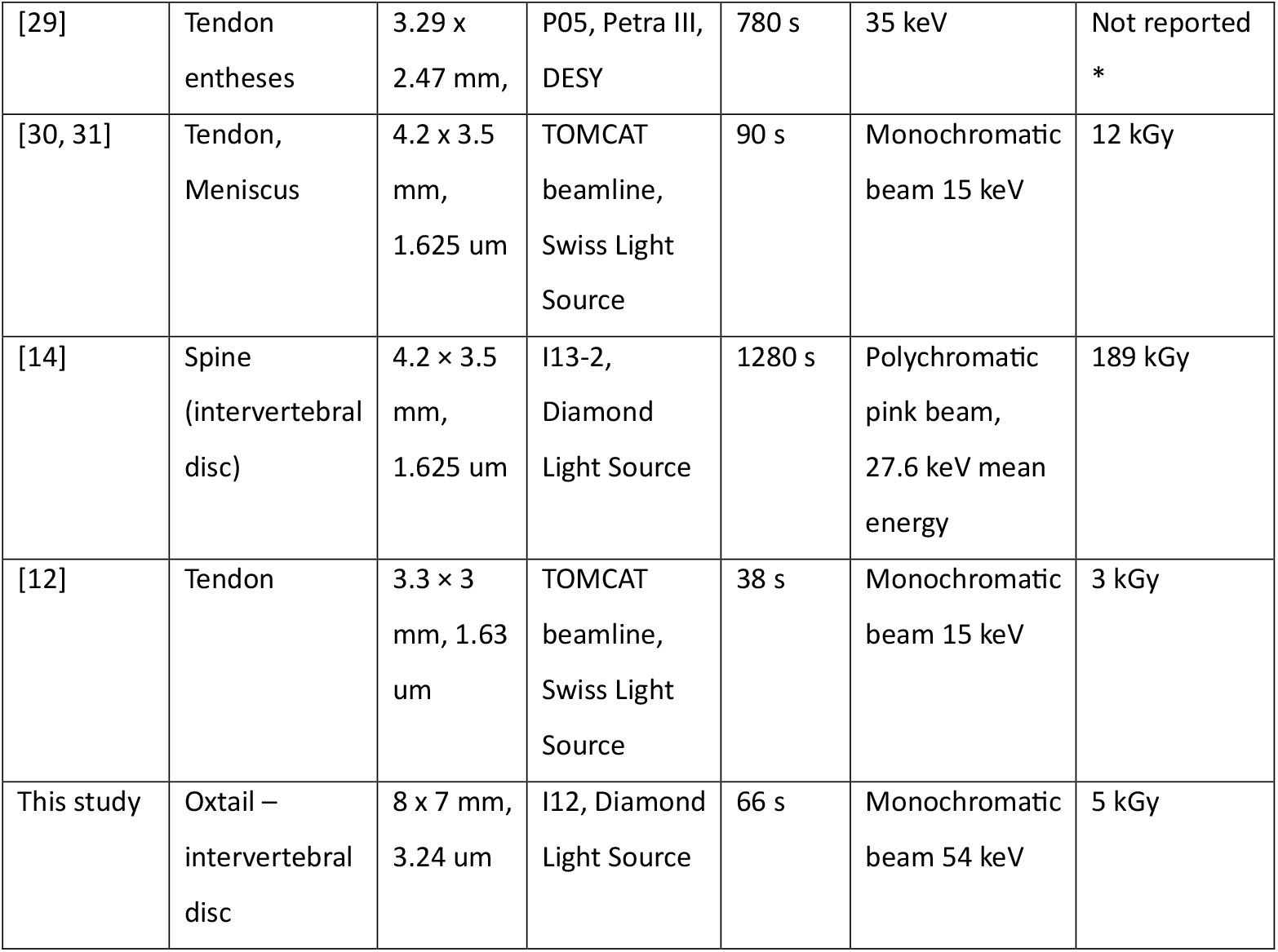
Recent musculoskeletal sCT in situ studies which mention dose. * damaged observed after several scans.

Reducing scan time is also critical for minimising radiation dose, particularly as the effects of synchrotron X-ray exposure on tissue microstructure and mechanical properties remain poorly understood—especially when multiple scans are required for *in situ* studies. Absorption-based imaging generates higher signal- and contrast-to-noise ratios compared to edge enhancement from Fresnel fringes used in propagation phase contrast imaging. Consequently, achieving shorter scan times for phase contrast imaging, which often requires summation of a lower signal-to-noise, can be more challenging than in absorption imaging.

In this study, we optimised imaging time to resolve the soft tissue by measuring image quality (using a perception-based image quality evaluator, PIQE) for a range of projection exposure times and number of projections. From the PIQE score plotted in Figure 2, scans less than 60 s had a rapid increase in PIQE score representing a decrease in image quality. A scan protocol using 0.018 s exposure time per projection and 2000 projections (with an additional 0.015 s overhead per projection) resulted in a total scan time of 66 seconds. This configuration provided sufficient image quality for analysis while maintaining a relatively low dose (~5 kGy per scan).

The dose per scan was comparable to the lowest reported in similar studies (Table 4). In general, higher-resolution imaging requires longer scan times and results in higher radiation doses. Compared to previous work, the parameters used here represent a favourable balance between field of view, resolution, scan time, and dose. After six scans, the total accumulated dose per sample was approximately 30 kGy—remaining below the commonly cited “safe” threshold of 35 kGy for bone tissue.

It may be possible to reduce scan times by using a rig with an open frame configuration, for example the rig used by Madi et al. [8]. In the current study, the Deben rig’s glassy carbon support tube had an absorption coefficient of 0.15 at 54 keV, which reduced overall X-ray transmission and attenuated the signal reaching the detector.

Shorter scan times can also be achieved by increasing frame rate. Frame rate can be improved by grouping pixel binning albeit with a trade-off in resolution [10] or with future innovations in high-speed camera acquisition. Additionally, developments in image reconstruction also have potential to reduce scan time. Iterative reconstruction algorithms have shown potential to enhance signal-to-noise from fewer projections [32]. More recently, deep learning-based denoising and reconstruction techniques have gained attention [33, 34], offering valuable tools for future low-dose imaging experiments.

### 4.2. Whole-sample mechanical behaviour was not affected by multiple scans

X-ray exposure has previously been associated with increased stiffness and brittle failure in bone tissue. However, in this study, the stiffness of the oxtail samples—measured from the linear region of the loading cycle—remained consistent across increasing levels of irradiation (Figure 4aii).

Additionally, the viscoelastic response of irradiated samples, assessed through hysteresis (Figure 4bii) and stress recovery time constants (Figure 4c), followed similar trends to those observed in unirradiated control samples across all loading cycle sets. These findings agree with the recent low dose in situ study by Pierantoni et al. [12] where tendon elastic modulus was unchanged.

Overall, variability between individual samples was greater than any variation observed across scan cycles, indicating that the differences between irradiated and unirradiated groups were too small to meaningfully affect the mechanical outcomes and favouring the use of *in situ* tomography to observe in real-time musculoskeletal samples behaviour under loading conditions. Future studies may consider applying higher cumulative doses to determine a threshold at which mechanical properties begin to deteriorate. However, given the practical constraints of *in situ* synchrotron computed tomography (sCT), six scans were sufficient to capture a meaningful time-lapse dataset without compromising tissue integrity.

### 4.2. Micro-scale changes were not observed in calcified tissue

No changes in microstructure or mechanical strain were observed in the calcified endplate with increasing numbers of scans. While previous studies have reported the development of strain and eventual microcracking in calcified tissues—even at relatively low doses (e.g. Fernández et al., 2018)—our findings contrast with these observations. In the present study, despite exposing samples to a comparable cumulative dose (~30 kGy over six scans), no microcracks or increases in strain were detected.

Importantly, strain values remained at the noise-floor level throughout the experiment, with median first and third principal strains consistently around 0.001 ε and no increase observed across scans (Figure 5d–e). The new results reported here are in contrast to an earlier study where Fernandez at al. [7] reported an increase in overall strain from below 0.001 at 9.4 kGy to 0.003 after 23.5 kGy.

Furthermore, less than 1% of values in Figure 5 were lower than the threshold criteria of 0.01 used by Fernandez et al. compared to 7% after 30 kGy and 23.5 kGy irradiation respectively. The differences between findings may be due to different sample types and experimental set up (Table 4).

We propose that the use of a higher-energy monochromatic beam (54 keV) in this study contributed to the reduced likelihood of radiation-induced damage. The photoionisation cross section, which reflects the probability of ionisation events, decreases as the incident photon energy increases relative to the binding energy of the target element. Calcium, the most abundant heavy element in the endplate, has a photoionisation cross section of 0.61 cm^2^/g at 54 keV, compared to 4.80 cm^2^/g at 29 keV—a 7.8-fold difference. This suggests that the lower-energy beam used in previous studies would have induced significantly more ionisation events.

Furthermore, other studies have used broader energy spectrum energy beams (Table 4) – such as laboratory sources or polychromatic ‘pink’ synchrotron beams - which, while increasing flux and reducing scan time, also include a greater proportion of low-energy photons. These lower-energy components contribute disproportionately to radiation damage. Beam filtering can mitigate this by removing the low-energy portion, but the use of a monochromatic beam, as in this study, inherently avoids this issue and is advantageous for reducing ionisation-related tissue damage.

### 4.4. Molecular changes were present in calcified tissues but not in soft tissues

Raman spectroscopy was used to assess molecular changes in the annulus fibrosus (AF), nucleus pulposus (NP), and bone following six scans, corresponding to a total radiation dose of 30 kGy. The AF and NP spectra exhibited minimal alterations, indicating that these soft tissues were largely unaffected by the radiation dose. However, small but statistically significant changes were observed in the AF spectra at 1318 cm^−1^ (CH_3_, CH_2_, proline) and 1340 cm^−1^ (glycosaminoglycans, δ C–H (CH_2_)), which may reflect variations in sample hydration or intrinsic biological variability. The NP spectra showed a slight shift in the Amide I band, potentially indicating subtle changes in protein secondary structure, such as alterations in α-helix or β-sheet content.

In contrast, the bone spectra revealed more pronounced changes, consistent with findings reported by Barth et al. Specifically, there was an increase in collagen-associated peaks (Amide I, CH_2_ bending mode, Amide III, and collagen backbone), indicative of enhanced collagen crosslinking. Additionally, irradiated bone samples exhibited a reduction in the phosphate peak and shifts in the carbonate substitution band, both of which are associated with bone mineralisation. These spectral changes provide further evidence that radiation exposure contributes to bone embrittlement, characterised by an increased elastic modulus due to collagen crosslinking and a concurrent reduction in mechanical strength resulting from mineral loss.

## 5. Conclusion

This study demonstrates the feasibility of low-dose synchrotron computed tomography (sCT) for high-resolution, *in situ* imaging of intact bovine intervertebral discs, while preserving tissue structure and mechanical integrity. By optimising scan parameters to achieve sufficient image quality within a 66-second acquisition time, we limited the total radiation dose to 30 kGy across six scans—remaining below the commonly cited safety threshold for bone tissue.

This work used a higher energy monochromatic X-ray beam (54 keV), which enabled adequate transmission through larger, hydrated soft tissue samples while reducing the likelihood of radiation-induced damage. Compared to lower energy beams commonly used in previous studies (~29 keV), the higher energy beam significantly reduced the photoionisation cross-section of calcium, the dominant attenuating element in mineralised tissues. This likely contributed to the absence of microstructural damage or strain accumulation in the calcified endplate, as confirmed by digital volume correlation (DVC) analysis.

Mechanical testing revealed no changes in stiffness, hysteresis, or stress recovery behaviour between irradiated and unirradiated samples, indicating that soft tissue mechanics were not adversely affected by repeated imaging. At the molecular level, Raman spectroscopy detected minimal changes in soft tissues. In contrast, bone spectra exhibited changes consistent with increased collagen crosslinking and reduced mineralisation, suggesting early signs of radiation-induced embrittlement.

Together, these findings support the use of high-energy, low-dose sCT for dynamic, multi-scan imaging of musculoskeletal tissues, particularly when combined with correlative mechanical and molecular assessments. Future work should explore the effect of different x-ray energies, photoionisation cross-sections, and dose thresholds for different tissue types, while leveraging advanced reconstruction and denoising techniques to further reduce exposure while maintaining image quality.

## Acknowledgments

The authors gratefully acknowledge facilities and research support provided by Diamond Light Source (beam time proposal MG33984) and the STFC Central Laser Facility (access proposal 23230023). C.M.D. is grateful for the support provided by the Wellcome Discovery Research Platform for Cell-Matrix Biology (226804/Z/22/Z).

## Abbreviations

(sCT): synchrotron CT
(DVC): digital volume correlation
(IVD): intervertebral disc
(AF): annulus fibrosus
(NP): nucleus pulposus

## Notes

### Competing Interest Statement

The authors have declared no competing interest.

