## Supplementary figures for "Preserving Tissue Integrity Under the Beam: High-Energy, Low-Dose Synchrotron CT for *in situ* Imaging of Bovine Intervertebral Discs"

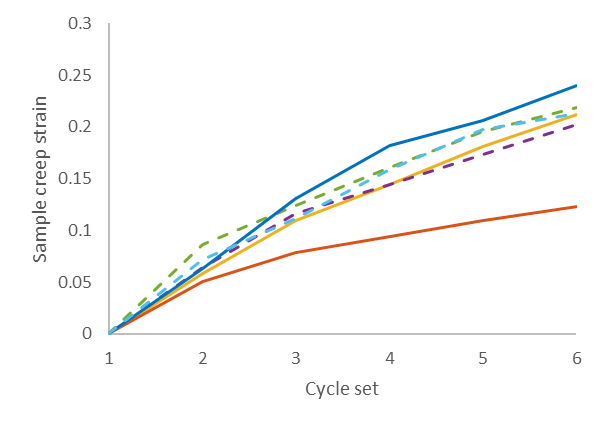

Supplementary figure 1 Sample creep strain after cyclic loading sets.

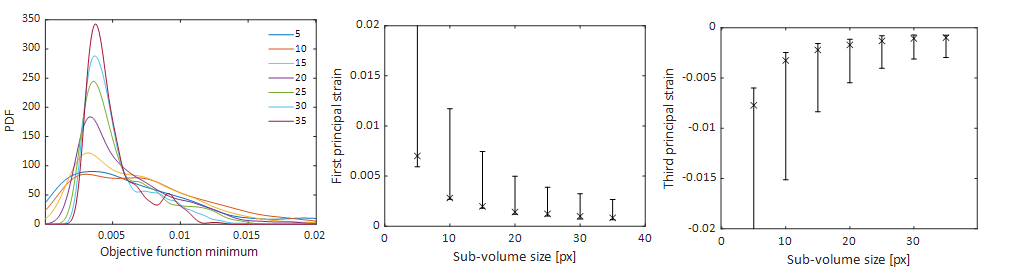

Supplementary figure 2 Optimising sub-volume size for DVC. a) Objective function (normalised sum squared difference) minimum. b) First principal strain. c) Third principal strain.

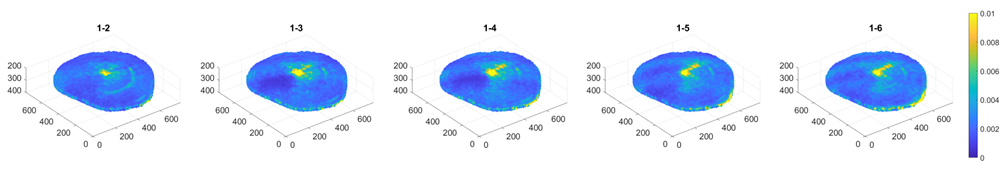

Supplementary figure 3 DVC objective function across all images showing centre-void and ring artefact patterns.
